# Diverse human astrocyte and microglial transcriptional responses to Alzheimer’s pathology

**DOI:** 10.1101/2021.07.19.452932

**Authors:** Amy M. Smith, Karen Davey, Stergios Tsartsalis, Combiz Khozoie, Nurun Fancy, See Swee Tang, Eirini Liaptsi, Maria Weinert, Aisling McGarry, Robert C. J. Muirhead, Steve Gentleman, David R. Owen, Paul M. Matthews

## Abstract

To better define roles that astrocytes and microglia play in Alzheimer’s disease (AD), we used single-nuclei RNA sequencing to comprehensively characterize transcriptomes in astrocyte and microglia nuclei isolated *post mortem* from neuropathologically-defined AD and control brains with a range of amyloid-beta and phospho-tau (pTau) pathology. Significant differences in glial gene expression (including AD risk genes expressed in astrocytes [*CLU, MEF2C, IQCK*] and microglia [*APOE, MS4A6A, PILRA*]) were correlated with tissue amyloid and pTau expression. Astrocytes were enriched for proteostatic, inflammatory and metal ion homeostasis pathways. Pathways for phagocytosis, proteostasis and autophagy were highly enriched in microglia and perivascular macrophages. Gene co-expression analyses revealed potential functional associations of soluble biomarkers of AD in astrocytes (*CLU*) and microglia (*GPNMB*). Our work highlights responses of both astrocytes and microglia for pathological protein clearance and inflammation, as well as glial transcriptional diversity in AD.

## Introduction

Astrocytes and microglia play important and potentially causal roles in the disease^1^. Activated astrocytes and microglia are found around amyloid plaques^2^ and genes associated with AD risk are enriched in both cell types, particularly in microglia^3,4^. However, the mechanisms by which microglia and astrocytes contribute to disease genesis, progression and response still are poorly defined; a recent meta-analysis suggested that the diversity of glial responses in late onset sporadic Alzheimer’s disease is not well captured by mouse models^5^.

Single-nuclei RNA-sequencing (snRNASeq) from *post mortem* brain tissue^6,7^ is transforming the potential to characterise the molecular neuropathology of AD at the level of single cells^8-10^. However, because of their lower cellular abundance in the brain, microglia and astrocytes have been poorly represented in most studies published to date (e.g., 449-3982 microglia, representing only 1-3% of the total nuclei annotated ^8,9,11^), limiting the depth to which they can be characterised. Astrocytes and microglia also have highly heterogeneous phenotypes^12-14^. To address these limitations, we employed a novel negative-selection approach that enriches for them in nuclei isolated from *post mortem* tissue. This allowed us to characterise snRNASeq transcriptomes from much larger numbers of these nuclei (52,706 astrocytes and 27,592 microglia) efficiently. After quantitative assessments of neuropathological features in each brain region, we generated transcriptional signatures associated with amyloid-beta or pTau pathology in non-diseased control (NDC) and AD brains. These data allowed us to develop comprehensive descriptions of gene co-expression networks that provide both further insights into responses of astrocytes and microglia to AD pathology and evidence for cell-type specific functions of genes associated with risk for AD. Transcription factors potentially responsible for the differential gene expression with pathology were identified from these co-expression modules. We confirmed our major observations by re-analyses of data from four previously reported AD snRNASeq studies. Our work provides new insights into linked glial-specific responses mediating pathological protein clearance and inflammation in AD, highlighting diverse roles of astrocytes and microglia.

## Results

### Selective astrocyte and microglia transcriptome sequencing

The proportions of microglia and astrocytes defined by snRNASeq of nuclei isolated from the human brain *post mortem* are low and variable^8,9^. To enable comprehensive analyses of differential transcript expression in astrocytes and microglia with AD pathology, we enriched for these glia by selectively removing neuronal (NeuN-positive^7^) and oligodendrocyte (Sox10-positive^15^) nuclei using FACS (Fig. 1 and Extended Data Fig. 1). For this, we isolated nuclei from each of two cortical regions (entorhinal and somatosensory cortex) taken from six brains with low levels of AD neuropathology provided by donors without reported cognitive impairment and from six brains with high levels of AD pathology (Extended Data Fig. 2). Astrocyte nuclei had a mean unique molecular identifier (UMI) count of 8,775 with an average of 3,166 distinct genes and microglial nuclei had a mean UMI count of 4,808 with an average of 2,132 genes. Amongst the 52,706 astrocyte nuclei, we found expression of 90% of astrocyte transcripts previously reported from human brain astrocytes (500 genes)^16^. 16/65 AD risk genes were represented in the astrocyte co-expression network (Extended Data Fig. 3a). The 27,592 total microglial nuclei included expression of 96% of the recently described microglial “core” consensus transcriptome (249 genes)^17^. Microglia also were highly enriched in genes associated previously with genetic risk for AD (27/65)^1,18,19^(Extended Data Fig. 4a).

**Figure 1:**
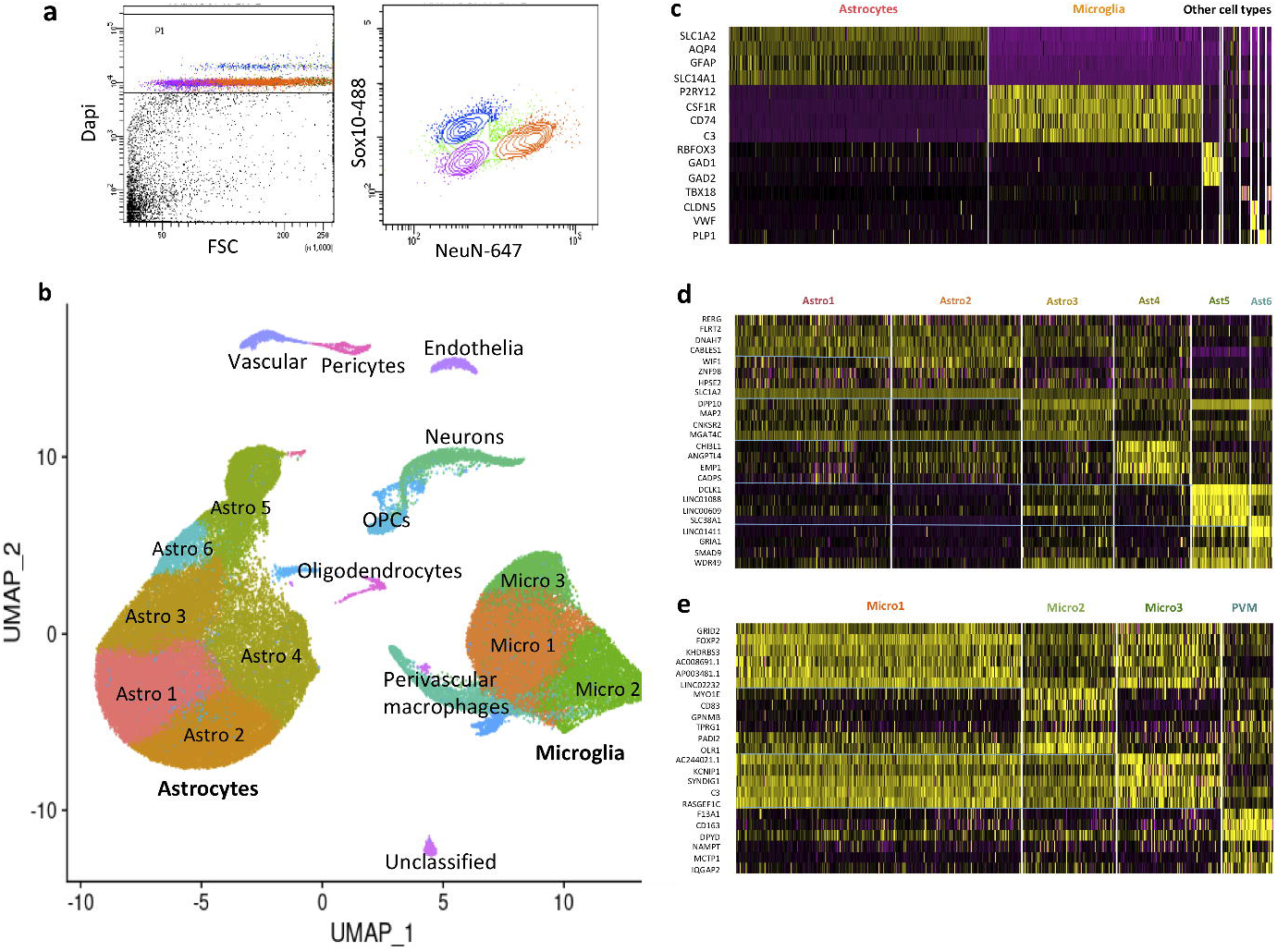
Analysis of human brain microglia and astrocytes from low and high AD pathology brains by single-nuclei RNASeq. a) FACS gating method for sorting of human brain nuclei to enrich for microglia and astrocyte double-negative population (pink; mean for 24 samples = 17.9%, standard deviation = 6.2%). Dapi+ve nuclei were selected first, followed by separation based on NeuN and Sox10 staining. b) UMAP 2D visualisation of clustering of 91,655 single nuclei from the 24 brain samples (average of 3,819 nuclei per sample) including 52,706 (58%) astrocytes and 27,592 (30%) microglia. Smaller numbers of nuclei from neurons, oligodendrocytes and vascular cells also were found (Extended Data Fig. 1), but these cell types formed distinct UMAP clusters that were not analysed further. c) Heatmap showing cell type-specific marker gene expression in the nuclei clusters. d) Heatmap of top differentially expressed genes in each microglial cluster. e) Heatmap of top differentially expressed genes in each astrocyte cluster.

### Increased expression of genes related to metal ion homeostasis, proteostasis and inflammation in astrocytes with AD pathology

Gene expression associated with extracellular amyloid plaques or intraneuronal neurofibrillary tangles (pTau) (Fig. 2) were discovered by regressing gene expression against amyloid-beta (expressed as log_2_ fold difference/% area stained) or pTau (expressed as log_2_ fold difference/% pTau positive cells) densities in sections prepared from homologous regions of the contralateral hemispheres for each of the brains. Half of the significantly positively associated genes expressed were correlated with both amyloid-beta and pTau pathology, but almost three-fold more transcripts were associated uniquely with amyloid-beta (313 genes) expression relative to pTau (106 genes) (Fig. 2). We found significant astroglial functional enrichment for pathways involved in the ‘cellular response to zinc ion’, ‘cellular response to copper ion’ and ‘response to metal ions’ with both amyloid-beta and pTau expression (Fig. 3); genes encoding proteins involved in metal ion homeostasis (*MT1G, MT1F, MT1E, MT2A, MT3* and *FTL*) were amongst the top transcripts most highly *positively* associated with pathology in astrocytes. Transcripts involved in ‘chaperone-mediated protein complex assembly’ and ‘response to unfolded protein’ pathways, such as *CRYAB, HSPB1, HSPH1* and *HSP90AA1* also were positively differentially expressed. Increased expression of the AD risk gene *CLU* was associated with pTau pathology in astrocytes (Extended Data Fig. 5). Expression of the AD risk gene *IQCK* was positively associated with both amyloid-beta and pTau (Extended Data Fig. 5). Pathways involved in inflammatory processes also were significantly enriched (‘NLRP3 inflammasome’ and ‘NFkB is activated and signals survival’). By contrast, “core” or homeostatic astrocyte transcripts, such as those for glutamate transporters *SLC1A3* and *SLC1A2* or for *IL-33* (CSF1R ligand, which promotes microglial synaptic remodelling^20^) were down-regulated. The AD risk-associated *MEF2C* transcription factor, as well as *MAFG, JUND, CEBPB, MAF* and *LHX2* were up-regulated, suggesting roles for these transcription factors in the regulation of responses to AD pathology.

**Figure 2:**
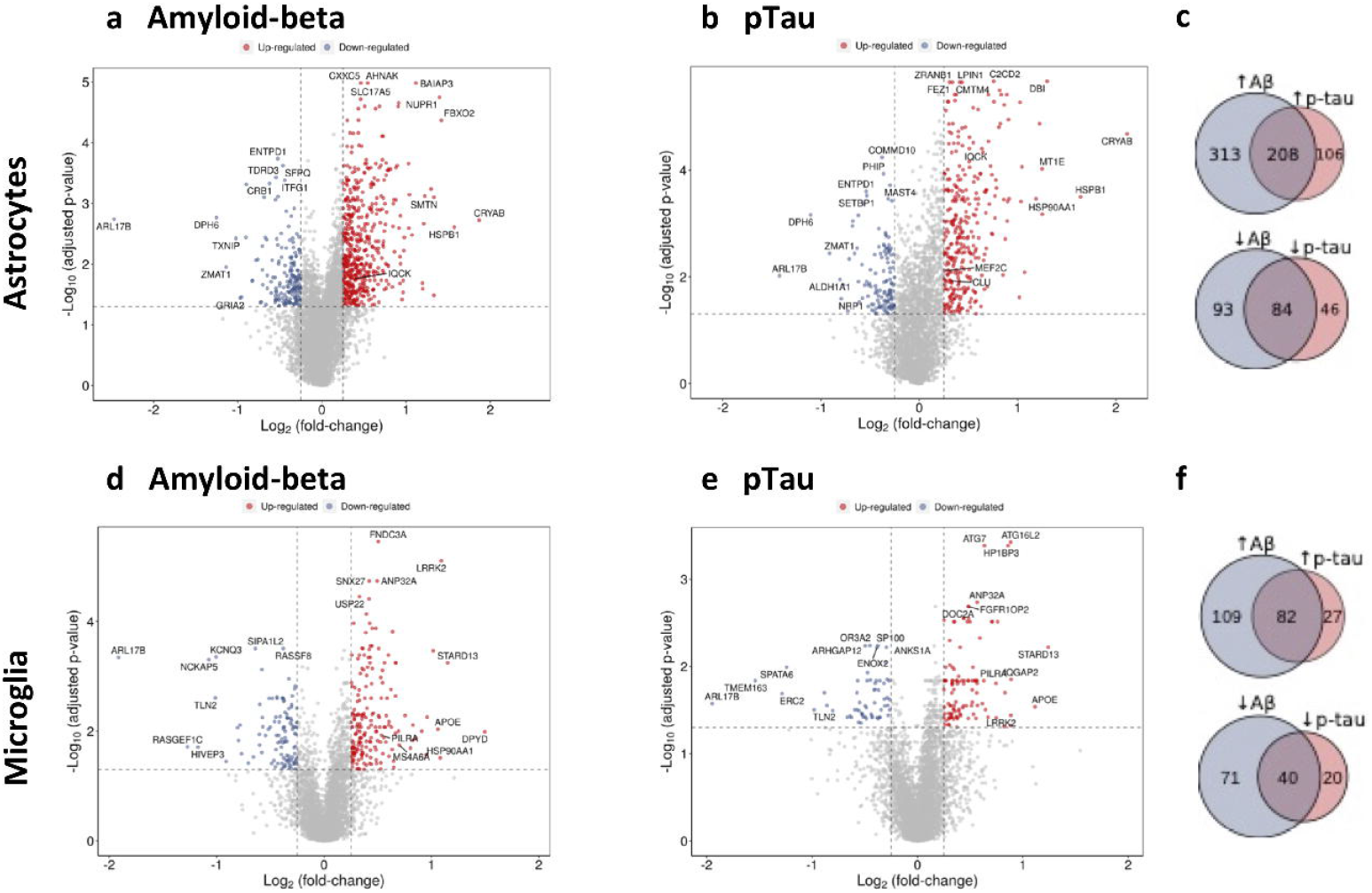
Differential gene expression in astrocytes and microglia with amyloid-beta and pTau pathology. a,b) Volcano plot of transcripts differentially expressed in astrocyte nuclei (threshold of ≤ 0.05 adjusted p-value, abs logFC≥ 0.25, omitting the top three most variable genes between samples) with immunohistochemically-defined tissue amyloid-beta (a) and pTau (b) density. c) Venn diagram illustrating the number of genes positively correlated (top) and negatively correlated (bottom) with amyloid-beta (blue) and pTau pathology (pink) in astrocytes. d,e) Volcano plot of transcripts differentially expressed in microglial nuclei (threshold of ≤ 0.05 adjusted p-value, abs logFC≥ 0.25, omitting the top three most variable genes between samples) with immunohistochemically-defined tissue amyloid-beta (d) and pTau (e) density. f) Venn diagram illustrating the number of genes positively correlated (top) and negatively correlated (bottom) with amyloid-beta (blue) and pTau (pink) pathology in microglia

**Figure 3:**
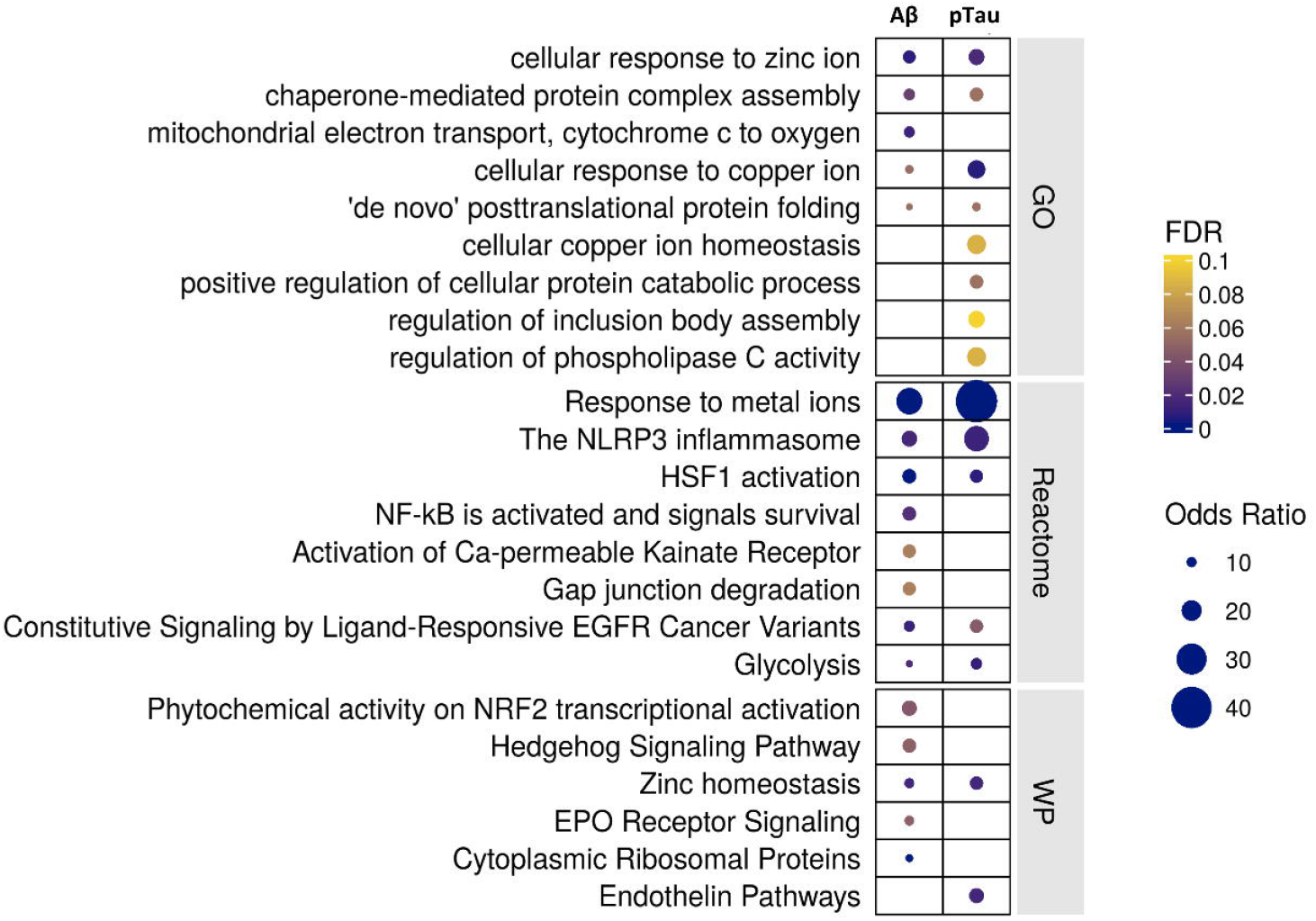
Functional enrichment of differential gene expression with amyloid-beta (left) and pTau (right) pathology in astrocytes. The plots describe the significant functionally enriched pathways in astrocytes obtained using enrichR (see Methods) from Gene Ontology (GO), Reactome and Wikipathways (WP) databases

We confirmed with gene set enrichment (AUCell) that transcripts positively differentially expressed with pathology also were significantly enriched in nuclei with human AD pathology reported in previously published snRNASeq studies ^8-10,21^, albeit with very low log fold changes in one^21^ out of the four datasets analysed.

### Expression of genes related to autophagy, phagocytosis and proteostasis in microglia with AD pathology

Microglial transcripts most highly *positively* associated with tissue amyloid-beta and tissue pTau density included those for genes associated with AD risk (*APOE, MS4A6A* and *PILRA*, Fig. 2 and Extended Data Fig. 5), as well as those for genes associated with risks for other neurodegenerative disorders (*LRRK2, SNCA* and *GPNMB*, associated with Parkinson’s disease^22^, and *GRN*, associated with ceroid lipofuscinosis^23^ and frontotemporal dementia^24^). Four-fold more transcripts were associated uniquely with amyloid-beta (109 genes) expression relative to pTau (27 genes), while 60% of the significantly positively associated genes expressed were correlated with both amyloid-beta and pTau pathology (Fig. 2). Differentially expressed transcripts were functionally enriched in ‘selective autophagy’ and ‘microglia pathogen phagocytosis’ pathways (Fig. 4); *ASAH1, ATG7, STARD13* and *MYO1E* were amongst the most strongly positively associated genes. Perivascular macrophages (PVM) also showed functional enrichment in ‘selective autophagy’, as well as several CCT/TriC molecular chaperone complex pathways involved in proteostasis and actin/tubulin folding (Extended Data Fig. 6). Up-regulation of the transcription factors *MAFG, MITF* and *JUND* with amyloid-beta or pTau pathology suggests their involvement in transcriptional regulation of these microglial and PVM responses to pathology.

**Figure 4:**
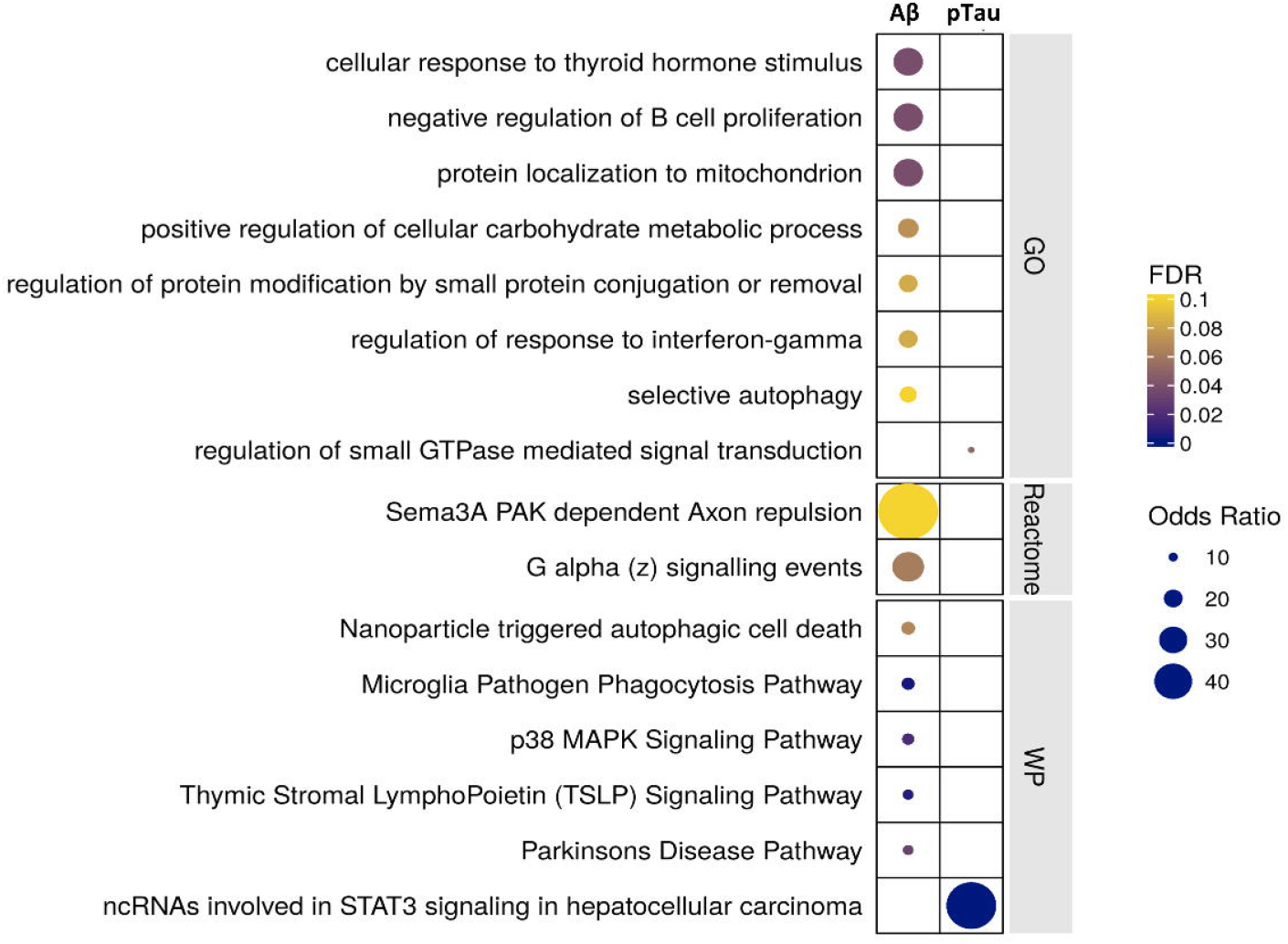
Functional enrichment of differential gene expression with amyloid-beta (left) and pTau (right) pathology in microglia. The plots describe the significant functionally enriched pathways in microglia obtained using enrichR (see Methods) from Gene Ontology (GO), Reactome and Wikipathways (WP) databases

We confirmed that transcripts positively differentially expressed in microglia were significantly enriched in nuclei with human AD pathology in previously published snRNASeq studies ^8-10,21^. Toll-like receptors (TLR2 and TLR10), HK2 (hexokinase 2), JAK2 (Janus kinase 2) and ITGAM (CD11b) were amongst the smaller number of transcripts in these cells that were significantly *negatively* associated with tissue amyloid-beta or pTau.

### Gene co-expression modules suggest glial cell-specific functional roles of AD GWAS genes

#### Evidence for involvement of CLU in astrocyte metal ion homeostasis and proteostasis pathways with AD

Co-expression network analyses were used to characterise gene expression modules (MEGENA) in astrocytes and microglia, suggesting potential functional relationships. AD GWAS genes *CLU* and *IQCK* were co-expressed in an astrocyte module (module 9) which was amongst the most strongly positively correlated with both amyloid-beta and pTau density. Both *CLU* and *GJA1* (Gap Junction Protein Alpha 1; Connexin-43) are hub genes in this module, which was functionally enriched in transcripts for proteins involved in metal ion homeostasis (e.g., ‘metallothioneins bind metals’ and ‘response to metal ions’) and proteostasis (‘HSF1 activation’, ‘response to unfolded protein’ and ‘chaperone-mediated protein complex assembly’). They also were hub genes in the related child module 30, which includes genes in pathways for ‘ceramide transport’ and ‘gap junction assembly’ (Extended Data Fig. 3b). *CLU* was associated with the astrocyte regulons (for transcription factors MAF, MAFG, JUND and CEBPB) that had the strongest correlations with pTau and amyloid-beta. *CLU*-containing modules and the associated regulons also were enriched in AD nuclei reported in earlier studies.

#### Evidence for a cell-specific role for APOE in microglia linking phagocytic, complement and inflammatory activation pathways in AD

*APOE*, the largest genetic risk factor for AD, was up-regulated in microglia with both pTau and amyloid-beta pathology. *APOE* was a hub gene in microglia co-expressed both with *TREM2* and inflammatory activation and response genes (e.g., *C1QB, C1QC, CD74, CTSB*) in a module functionally enriched for pathways including the ‘endosomal/vacuolar pathway’, ‘microglia pathogen phagocytosis’, and ‘antigen processing – cross presentation’ (module 19) (Extended Data Fig. 4b). Regulons inferred to be responsible for microglial *APOE* expression included those for transcription factors MXD4, MITF, PBX3 and JUND. *APOE* expression in astrocytes was not significantly correlated with either amyloid or pTau pathology and co-expression relationships suggested a different functional role for *APOE* in astrocytes as a hub gene in a module functionally enriched for ‘dermatan sulphate biosynthesis’, ‘extracellular matrix organisation’ and ‘ferroptosis’ (module 13).

#### Microglial and PVM GPNMB are up-regulated with AD pathology in modules related to lipid homeostasis

Glycoprotein nonmetastatic melanoma protein B (GPNMB) is elevated in plasma and CSF with AD and has been proposed as a biomarker of disease^25^. We found that *GPNMB* is up-regulated in microglia with amyloid-beta and pTau pathology and in PVMs with pTau pathology. Consistent with this, *GPNMB* was found in association with the MAFG, JUND, MAFB, CEBPD and CEBPA transcription factor regulons which showed strong correlations to amyloid-beta and pTau in microglia (Fig. 5a). *GPNMB* also is a hub gene in microglial co-expression modules 11 and 34, which are strongly associated with amyloid-beta and pTau expression. As well as *GPNMB*, hub genes for module 11 also include *ASAH1, ATG7, STARD13, IQGAP2, CPVL, TANC2* and *MITF*, all of which were positively differentially expressed with one or both of the AD pathologies (Fig. 5a). We found that module 11 was enriched in AD (relative to control) samples from three out of four previous human snRNASeq studies analysed^8,10,21^. Pathways involving the differentially expressed genes in module 11 suggest a functional role in cholesterol homeostasis (‘regulation of cholesterol storage’). We found that the smaller module 34 was enriched in AD samples from two out of four previous studies^9,21^. Functional pathways enriched in the *GPNMB* hub gene module 34 relate to phospholipid and lipoprotein homeostasis (‘phospholipid efflux’, ‘phospholipid homeostasis’, ‘lipoprotein metabolism’).

**Figure 5:**
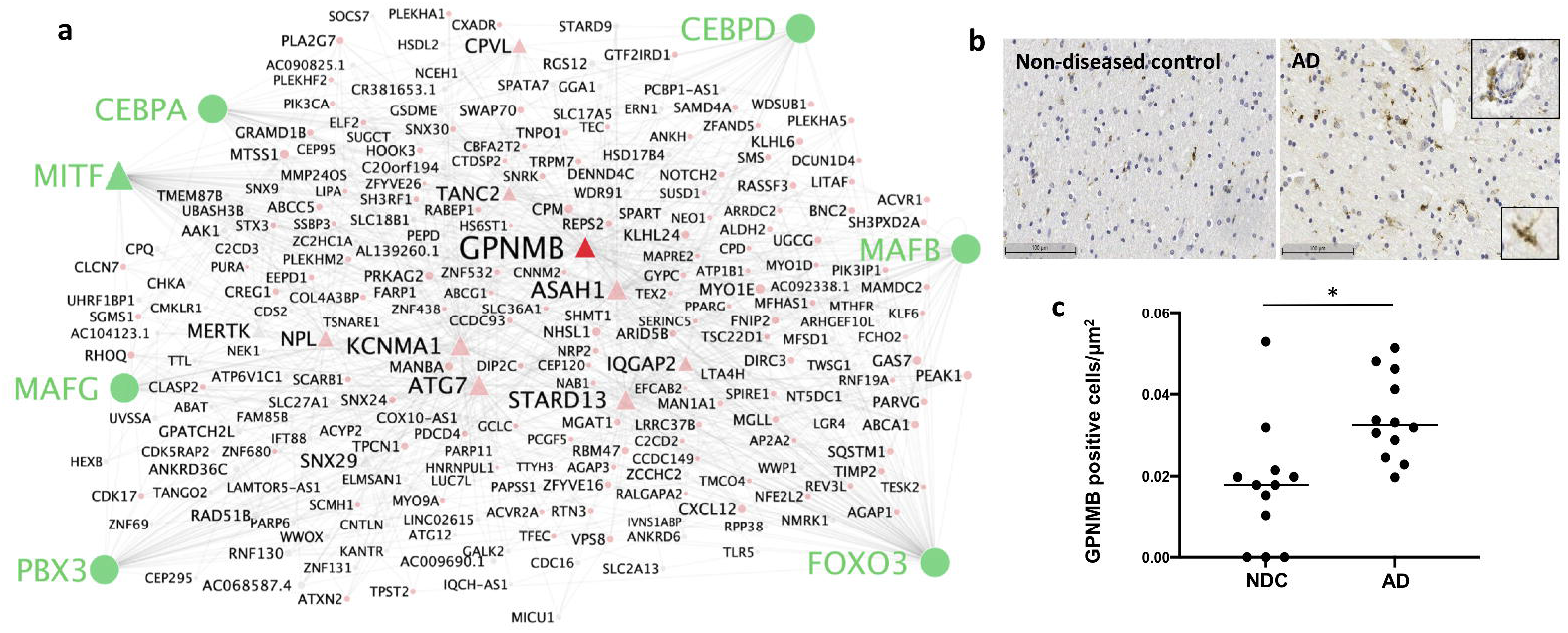
*GPNMB* is a hub gene in microglial gene co-expression modules up-regulated with AD pathology. a) Graph of microglial gene co-expression module 11 enriched with amyloid-beta and pTau pathology for which *GPNMB* is a hub gene (Triangles = module hub genes). *GPNMB* is expressed in regulons identified by their transcription factors (in green). b) Immunohistochemical staining of GPNMB in *post mortem* human brain tissue from representative NDC and AD brains with nuclear counterstain. Staining is present in both microglia and perivascular macrophages (insets). Scale = 100μm. c) Quantification of density of GPNMB-positive cells in cortical tissue by automated image analysis showing an increase in GPNMB-positive cells per μm^2^ for AD cases compared to NDC (p = 0.0023; Mann Whitney test, two-tailed; N = 12). Each point represents a single sample and the horizontal bar indicates the median.

Consistent with the expected GPNMB expression in PVM, immunohistochemistry for GPNMB showed prominent staining around blood vessels, as well as in a subset of parenchymal microglia (Fig. 5b). There was a mean ∼1.8-fold increase in GPNMB-positive cells/μm^2^ in AD (relative to NDC) in the cortical tissue studied here (Fig. 5c). GPNMB staining density in sections of samples was positively correlated both with tissue pTau (R=0.39) and with expression of modules 11 and 34 (R=0.507 and 0.576, respectively) defined by snRNASeq of the same region in the contralateral hemisphere.

### Transcriptional heterogeneity associated with AD pathology in sub-sets of human astrocytes and microglia

Our snRNASeq data reduction identified six distinguishable clusters that expressed the core set of astrocyte genes. Each also expressed distinct sets of genes that suggested sub-types of astrocytes (Fig. 1). Astro1 and Astro2 expressed higher levels of genes involved in core astrocyte functions, such as *SLC1A2* (*GLT1*) and glutamine synthetase (*GLUL*) and Astro2 was enriched for pathways including ‘neurotransmitter uptake’, ‘glutamatergic synapses’ and ‘amino acid import’. In contrast, Astro4 and Astro5 were characterized by relative expression of genes involved in extracellular matrix formation and functions^26^ and were enriched for pathways including ‘carbohydrate binding’ and ‘cell-matrix adhesion’. Astro4 was distinguished from Astro5 by relatively high expression of *VEGFA*, while Astro5 was enriched uniquely for immune response pathways, such as the Toll-like receptor cascade and the activated astrocyte marker *GFAP*. Transcripts in Astro6 were enriched for metallothionein genes. Astro5 and Astro6 showed the greatest specificity for regulons most highly up-regulated with pTau (for transcription factors JUND, MAF and CEBPB).

The disease associated astrocyte (DAA) expression signature defined in a mouse model^27^ was represented in all astrocyte clusters other than Astro3. 4.6% of DAA genes were up-regulated with amyloid-beta and 9.2% with pTau. Although the A1 (12/15 genes expressed) or A2 (13/13 genes expressed) gene sets^28^ associated previously with injury-responsive or homeostatic astrocytes in rodent models, respectively, were represented in the astrocyte clusters, neither A1 nor A2 gene sets were enriched significantly in the total astrocyte nuclei or in any of the clusters.

Clustering parameters selected to distinguish PVM (identified by markers such as *CD163, MRC1* (*CD206*) and *MSR1*^29^) also identified three clusters of nuclei (designated Micro1, 2 and 3) expressing different sets of microglial marker genes (Fig. 1). Micro1 was most highly enriched in transcripts for human ‘core’^30^ homeostatic genes, Micro2 showed a relative functional enrichment for the ‘TYROBP causal network’ and ‘ferroptosis’ pathways and Micro3 expressed lower levels of both homeostatic and activation genes, but had higher expression of *C3* and *LPAR6*. Micro2 showed the greatest specificity for the MAFG, CEBPA, JUND, CEBPD and MITF regulons found here to be highly correlated with amyloid-beta and pTau. Plaque-induced (PIG^31^), disease associated (DAM^32^), activated response (ARM^33^) and interferon response microglial (IRM^33.34^) gene sets identified in rodent amyloid models were expressed in all of the clusters, with relative enrichment of DAM, ARM and, to a lesser degree, IRM gene sets in Micro2 (Extended Data Fig. 7). PVMs were also relatively enriched in these gene sets related to microglial activation, as well as in a gene set associated with human microglial aging^35^ (Extended Data Fig. 7).

## Discussion

Our results describe potentially neuroprotective gene up-regulation for proteostasis, phagocytosis and protein clearance with amyloid-beta and pTau pathology in both astrocytes and microglia. A 3-4 fold greater number of genes were uniquely differentially expressed with amyloid-beta relative to pTau, mirroring observations with microglia in preclinical transgenic models overexpressing the two proteins^36^. Differentially expressed genes in astrocytes were associated with increased expression of genes for the regulation of metal ion homeostasis that may both protect cells from oxidative injury and facilitate redox-dependent chaperone clearance of abnormal proteins by astrocytes^37^. However, astrocytes also expressed inflammasomes and inflammatory activation pathways commonly found in microglia, both of which may promote neurodegeneration. Transcriptional functional diversity suggests distinguishable sub-types of human astrocytes and microglia enriched in gene sets related to (but distinct from) those previously described in transgenic amyloid or tau mouse models or with aging^27,32,33,35^. Moreover, co-expression networks suggest cell-specific functional roles for genes associated with AD risk, most notably highlighting *APOE* as a hub gene in microglial co-expression modules linking gene expression subserving phagocytic, complement and inflammatory activation pathways.

We extended previous approaches to enrich our *post mortem* brain snRNASeq data for microglia and astrocytes in our study. Several recent publications have described approaches for selective glial nuclei isolation from biopsy and rapid *post mortem* delay tissue, including the use of antibodies to microglial transcription factor PU.1^38^, a combination of NeuN and Olig2^39^, and IRF8^40^. Our negative-selection approach is distinguished by relying on nuclear markers that we have found to be robust to *post mortem* delay, reliably at least to 24 hr. Our analytical approach for addressing heterogeneity in the samples also differs from most prior studies (although it is not without similar precedents^11,41^). Categorical descriptions of brains by Braak stage do not directly reflect tissue pathology in the local regions studied, so we associated each glial nuclear preparation with quantitative measures of pTau and amyloid-beta density specific to the regions studied in each of the brains. We also used of a mixed regression model to account for uncontrolled sources of variance related to each sample^42^.

Astrocytes expressed genes suggesting functional roles for proteostasis with amyloid-beta and pTau-associated functional enrichment for HSF1 activation and chaperone pathways. This was accompanied by expression of genes for pro-inflammatory NF-kB and inflammasome pathways. Greater regional amyloid-beta and pTau also was associated with enrichment of astrocytes for metal ion homeostasis pathways and the expression of metallothioneins ^43-45^. Identification of *CLU* as a hub gene in astrocyte co-expression modules including genes for metal ion homeostasis and proteostasis may reflect dependence of clusterin chaperone functions on the local redox environment^37^. We found increased expression of the astrocyte *CLU* module with greater amyloid or pTau.

Co-expression analysis in microglia identified *APOE* as a hub gene for a module including complement genes (*C1QA, C1QB, C1QC*) and functionally enriched in phagocytosis pathways. Microglia may contribute to astrocyte activation through C1q expression and C1q and other complement cascade proteins co-localise with amyloid plaques in AD^46^. These analyses also highlighted potential functional relationships of *GPNMB* (concentrations of the protein product of which are increased in brain samples and cerebrospinal fluid of sporadic AD patients^25^) with a diverse group of genes connected to transcription factors whose regulons were up-regulated with amyloid and pTau, suggesting that GPNMB provides a biomarker for a human microglial neurodegenerative activation state central to pathological responses in AD.

We tested the relevance of microglial sub-cluster expression signatures defined in mouse transgenic models of AD using gene set enrichment analyses (Extended Data Fig. 7). We found evidence for similar, moderate levels of relative enrichment for inflammatory activation gene sets (DAM and ARM) in Micro2 and PVM^32,33^. The relatively low expression of both homeostatic and activation genes in Micro3 corresponded to patterns associated with ‘dystrophic’, immuno-senescent microglia^47^. Transcripts from the disease associated astrocyte (DAA) gene set were up-regulated with amyloid-beta and pTau, but not restricted to any particular astrocyte sub-set^27^. Our results highlight distinct features and a greater functional diversity amongst microglia and astrocytes in the human disease relative to related preclinical models.

We recognise limitations of our data, as well as their considerable promise for further exploration. Although restricted to end-stage tissue *post mortem*, we attempted to maximise the dynamic range of pathology sampled by evaluating correlations with quantitative pathological measures across two anatomical regions and brains of different Braak stages. We tried to compensate for the sparse 10X technology sampling of the transcriptome by increasing the number of glial nuclei of interest and provided evidence for generalisability of our results by demonstration that major co-expression modules, regulons and gene expression analyses discovered here also were represented in previous human *post mortem* snRNASeq datasets^8-10,21^. Even so, the nuclear transcriptome may be biased relative to that from a whole cell, potentially reducing the power to detect some genes reported from studies of related pathologies in mouse models^48^. The need to maximise detection sensitivity motivated us to enrich our sample for microglia and astrocytes for maximising the number of nuclei characterised and to make use of co-expression-based analyses, which rely less on detection of absolute expression levels than do single gene differential expression analyses.

Our work has extended previous studies defining AD-associated molecular pathology of glial cells substantively by describing proteostasis, metal ion homeostasis and inflammatory mechanisms in astrocytes and phagocytotic, proteostatic and autophagic pathways in microglia. We found that gene sets described in transgenic mouse models are variably represented in AD in the context of more complex glial phenotypes. Our data add to evidence that there are functionally distinct sub-types of astrocytes and microglia, with particular diversity amongst the former, in which we distinguished enrichments for gene sets associated with synaptic function, extracellular matrix formation, immune responses and control of metal ion homeostasis/redox state. While this diversity suggests multiple potential targets for therapeutic modulation, the complexity of human astroglia and microglial phenotypes simultaneously expressed in AD also needs to be taken into account.

## Methods

### Brain tissue

This study was carried out in accordance with the Regional Ethics Committee and Imperial College Use of Human Tissue guidelines. Cases were selected from the London Neurodegeneration (King’s College London) and Parkinson’s UK (Imperial College London) Brain Banks. Entorhinal and somatosensory cortex from 6 non-diseased control (NDC) cases (Braak stage 0-II) and 6 AD cases (Braak stage III-VI) were used (total of 24 brain samples). Cortical samples from two regions were prepared from each brain in order to characterise transcript expression with both higher (entorhinal cortex) and lower (somatosensory cortex) tissue densities of pTau in neurofibrillary tangles and amyloid-beta plaques. Brains used for this study excluded cases with clinical or pathological evidence for small vessel disease, stroke, cerebral amyloid angiopathy, diabetes, Lewy body pathology (TDP-43), or other neurological diseases. Where the information was available, cases were selected for a brain pH greater than 6 and all but one had a *post mortem* delay of less than 24 hr.

### Immunohistochemistry

Immunohistochemical staining was performed on formalin-fixed paraffin-embedded sections from homologous regions of each brain used for snRNASeq. Standard immunohistochemical procedures were followed using the ImmPRESS Polymer (Vector Laboratories) and SuperSensitive Polymer-HRP (Biogenex) kits (Table 2). Briefly, endogenous peroxidase activity and non-specific binding was blocked with 0.3% H_2_O_2_ and 10% normal horse serum, respectively. Primary antibodies were incubated overnight at 4°C. Species-specific ImmPRESS or SuperSensitive kits and DAB were used for antibody visualization. Counter-staining for nuclei was performed by incubating tissue sections in haematoxylin (TCS Biosciences) for 2 min. AD pathology was assessed by amyloid plaque (4G8, BioLegend 17-24) and pTau (AT8, NBS Biologics) staining. GPNMB staining in microglia was assessed using R&D antibody AF2550.

### Quantitative image analysis

Labelled tissue sections were imaged using a Leica Aperio AT2 Brightfield Scanner (Leica Biosystems). Images were analysed using HALO software (Indica Labs, Version 2.3.2989.34). The following image analysis macros were used for the study: area quantification macro (amyloid), multiplex macro (pTau) and microglia macro (GPNMB). GraphPad Prism version 8 for Windows (GraphPad Software, La Jolla, CA, USA) was used for plotting immunohistochemical results and performing statistical analysis. A Mann Whitney test was used to test for the significance of differences between NDC and AD.

We found that densities of pTau staining were greater in brains from the higher Braak stages, and that pTau and amyloid pathology both were found in the regions studied even in brains with lower Braak stages (Extended Data Fig. 2). This variability of local tissue pathology within similar Braak stages emphasises the potential importance of matching regional neuropathology with transcript expression for each brain individually.

### Nuclei isolation and selective glial enrichment

Fresh frozen entorhinal and somatosensory cortical tissue blocks were sectioned to 80 μm on a cryostat and grey matter separated by scoring the tissue with sharp forceps to collect ∼200 mg grey matter in an RNAse-free Eppendorf tube. Nuclei from NDC and AD samples were isolated in parallel using a protocol based on Krishnaswami *et al*. (2016)^6^. All steps were carried out on ice or at 4°C. Tissue was homogenized in a 2 ml glass douncer containing homogenization buffer (0.1% Triton-X + 0.4 U/μl RNAseIn + 0.2 U/μl SUPERaseIn). The tissue homogenate was centrifuged at 1000 g for 8 min, and the majority of supernatant removed without disturbing the tissue pellet. Homogenised tissue was filtered through a 70 μm filter and centrifuged in an Optiprep (Sigma) density gradient at 13,000 g for 40 min to remove myelin and cellular debris. The nuclei pellet was washed and filtered twice in PBS buffer (PBS + 1% BSA + 0.2 U/ml RNAseIn). Isolated nuclei were labelled in suspension in 1 ml PBS buffer with 1:500 anti-NeuN antibody (Millipore, MAB377, mouse) and 1:250 anti-Sox10 antibody (R&D, AF2864, goat) for 1 hr on ice. Nuclei were washed twice with PBS buffer and centrifuged at 500 g for 5 min. Nuclei were incubated with Alexa-fluor secondary antibodies at 1:1000 (goat-anti-mouse-647 and donkey-anti-goat-488) and Dapi (1:1000) for 30 min on ice, and washed twice. Nuclei were FACS-sorted on a BD Aria II, using BD FACSDiva software, gating first for Dapi +ve nuclei, then singlets and then Sox10- and NeuN-negative nuclei. A minimum of 150,000 double-negative nuclei were collected.

### Single nucleus capture and snRNA sequencing

Sorted nuclei were centrifuged at 500 g, resuspended in 50 μl PBS buffer and counted on a LUNA-FL Dual Fluorescence Cell Counter (Logos Biosystems, L20001) using Acridine Orange dye to stain nuclei. Sufficient nuclei were added for a target of 7,000 nuclei for each library prepared. Barcoding, cDNA synthesis and library preparation were performed using 10X Genomics Single Cell 3’ Gene Expression kit v3 with 8 cycles of cDNA amplification, after which up to 25 ng of cDNA was taken through to the fragmentation reaction and a final indexing PCR was carried out to 14 cycles. cDNA concentrations were measured using Qubit dsDNA HS Assay Kit (ThermoFisher, Q32851), and cDNA and library preparations were assessed using the Bioanalyzer High-Sensitivity DNA Kit (Agilent, 5067-4627). Samples were pooled to equimolar concentrations and the pool sequenced across 24 lanes of an Illumina HiSeq 4000 according to the standard 10X Genomics protocol. The snRNAseq data will be made available for download from the Gene Expression Omnibus (GEO) database (https://www.ncbi.nlm.nih.gov/geo/) under accession number GSE160936.

### Processing of FASTQ files, dimensionality reduction and clustering

snRNASeq data were pre-processed and clustered using 10X Genomics Cell Ranger and Seurat analysis tools^10,39,49^. Illumina sequencing files were aligned to the genomic sequence (introns and exons) using GRCh38 annotation in Cell Ranger v3.1. Nuclei were identified above background by the Cell Ranger software. Filtered gene matrices from CellRanger were loaded into R where Seurat v3 single-cell analysis package was used for analysis^50^. Genes that were expressed in three or more nuclei were used for further analysis. Further QC was performed to exclude nuclei with less than 200 genes or greater than 6000 genes or 25,000 UMIs, which likely represent low quality or doublet nuclei, respectively. Nuclei with greater than 5% mitochondrial genes were also excluded. Mitochondrial gene reads were excluded. The 24 samples had an average of 3819 nuclei per sample after passing QC.

Data was normalized and scaled using the *NormalizeData* function with normalisation.method□=□”LogNormalize” and scale.factor□=□10,000. Variable genes then were identified using the *FindVariableFeatures* function with nfeatures (number of variable genes) set to 2000. To integrate the data from all samples, *FindIntegrationAnchors* (dims = 1:20) and *IntegrateData* (dims = 1:20) functions were used. PCA analysis was run, using variable genes, for the top 30 components. Clusters were identified using *FindClusters* (resolution = 0.5) and UMAP was used for 2D visualization of clusters (with the top 15 PCs, based on the ranking of PCs by the variance explained by each).

Cell-type identification of clusters was performed using AUCell (see below) with cell-type specific genes identified by previous human brain snRNASeq studies ^51^. Cell type annotation was confirmed by visual inspection of key marker genes (Fig. 1 and Extended Data Fig. 1). Glial cluster specific genes were identified using the *FindMarkers* function. All clusters were composed of nuclei from all samples and did not represent a single case or disease group. A small number of nuclei that either did not express any major cell-type markers, or expressed a combination of cell-type markers, were categorized as ‘unclassified’. We used the thresholding method described above for identification and removal of unclassified clusters (Extended Data Fig. 1).

### Differential gene expression analysis

MAST was used to identify genes differentially expressed (associated) with histopathological features (using pTau or amyloid-beta as markers)^42,52^ to perform zero-inflated regression analysis by fitting a mixed model. The model specification was zlm(∼histopath_marker + (1|sample) + cngeneson + pc_mito + sex + brain_region, sca, method = “glmer”, ebayes = F). The fixed effect term pc_mito accounts for the percentage of counts mapping to mitochondrial genes. The term cngeneson is the cellular detection rate. Each nuclei preparation was considered as a distinct sample for the mixed effect. Models were fit with and without the dependent variable and compared using a likelihood ratio test. Units for differential expression are defined as log_2_ fold difference/% pTau positive cells (or log_2_ fold difference/% amyloid area), i.e., a one unit change in immunohistochemically-defined pTau (or amyloid density) is associated with one log_2_ fold change in gene expression. Genes expressed in at least 10% of nuclei from each cell type (either total microglia or total astrocytes) were tested. Genes with a log_2_ fold-change of at least 0.25 and adjusted p-value <0.05 were defined as meaningfully differentially expressed. As an additional filter, the percentage of the inter-individual variance in expression between the NDC subjects was calculated for each gene and three genes with unusually high (>2 standard deviations) variance (*LINGO1, SLC26A3* and *RASGEF1B*) in one or two samples were excluded.

### Gene ontology and pathway enrichment analysis

The gene ontology (GO) enrichment and the pathway enrichments analysis were carried out using the R package enrichR (v 3.0), which uses Fisher’s exact test (Benjamini-Hochberg FDR < 0.1)^53,54^. Genesets with minimum and maximum genes of 10 and 500 respectively were considered. To improve biological interpretation of functionally related gene ontology and pathway terms and to reduce the number of redundant gene sets, we first calculated a pairwise distance matrix using Cohen’s kappa statistics based on the overlapping genes between the enriched terms and then performed hierarchical clustering of the enriched terms^55^.

### Gene Set Enrichment Analyses (GSEA)

AUCell^56^ (R package v1.6.1) was used to quantify the expression of published gene set signatures. Mouse genes were converted to human orthologues where applicable using bioDBnet. Normalised data was processed in AUCell using the *AUCell_buildRankings* function. The resulting rankings, along with the gene lists of interest, were then put into the function *AUCell_calcAUC* (aucMaxRank set to 1% of the number of input genes). Resulting AUC scores were scaled across clusters and plotted in a heatmap (Extended Data Fig. 7).

We also used AUCell to test for enrichment of the gene sets that we identified on previously published human single-nuclei data ^8-10,21^. Where possible for the AUCell tests for enrichment of gene sets from previously reported data, the cell type annotations from the published data were used. Filtered matrices were processed and cell types identified using the methods described above for the Zhou *et al*.^*10*^ data set. The scFlow single cell analysis pipeline (https://nf-co.re/scflow) was employed for analysis of the Gerrits *et al*. data set. In brief, quality control was performed using the same criteria as described above. Sample integration was performed using Liger^57^ and dimension reduction using PCA and UMAP and finally clustering was performed using the Leiden algorithm^58^. Microglia and astrocytes were then identified using the same sets of marker genes used for the primary analysis (Fig. 1). AUCell was then run separately on microglia and astrocyte populations from each study using lists of our significantly up-regulated and down regulated genes with pTau and amyloid pathology, from microglia and astrocytes (thresholded as described above). The aucMaxRank term was set to 200 genes. LogFC values between Control and AD samples were estimated using the limma package in R^59^, using the default configuration and the following linear model: ∼diagnosis+nFeature, where nFeature is the total number of distinct features expressed in each nucleus (to account for the fact that nuclei that express a higher number of features may have higher AUCell scores).

Over-representation analyses of literature gene sets in our glial sub-clusters and differentially expressed gene lists were performed by Fisher’s exact test using the “enrichment” function of the R package “bc3net” (https://github.com/cran/bc3net). The p values associated with the Fisher’s exact test correspond to the probability that the overlap between the literature gene sets and the sub-cluster markers/differentially regressed genes from our dataset has occurred by chance.

### Gene co-expression (module), regulatory network (regulon) and enrichment analyses

Gene co-expression module and hub-gene identification analysis were performed separately for microglial and astrocyte populations using the MEGENA (v1.3.7) package^59^. The top 15% most variably expressed genes were used as input^13^. We evaluated the mean expression of each of these genes across all the nuclei in the expression matrices: all the genes in both astrocyte and microglia matrices were among the top 25% most highly expressed genes confirming that the choice to filter the expression matrix based on variability did not bias the inclusion towards genes with a particularly low level of expression. In addition, we verified that the filtered expression matrices for both astrocytes and microglia included a substantial proportion of the differentially expressed genes: over 90% for astrocytes and all but two of the microglial differentially expressed genes were included in the respective filtered expression matrices, suggesting that the filtered expression matrices contained a biologically meaningful gene set. The MEGENA pipeline then was applied using default parameters, using Pearson’s correlations and a minimum module size of 10 genes. Parent modules were produced from which a sub-set of genes form smaller child modules. For downstream analysis, interpretation and presentation of results, modules with >20 genes were retained. Co-expression modules were represented graphically using Cytoscape software (Mac OS version 3.8.0)^60^ with hub genes represented with a triangle and nodes with a circle with a diameter proportional to the node degree^59^. Genes previously associated with AD as defined by Kunkle *et al*. (Table 1^19^, 23 genes), Jansen *et al*. (Table 1^18^, a further 17 genes) and Andrews *et al*. (Table 1^1^, a further 25 genes), for a total of 65 genes, were annotated in the co-expression networks.

**Table 1:** Cohort information

|  | **M:F ratio** | **Age at death (yrs, mean +/- SD)** | **Post mortem delay (hr, mean +/- SD)** | **RIN (mean +/- SD)** |
| --- | --- | --- | --- | --- |
| **Non-diseased controls (Braak 0-II)** | 4:2 | 79.3 +/- 6.5 | 18 +/- 6.9 | 4.9 +/- 2.0 |
| **Alzheimer's disease (Braak III-VI)** | 4:2 | 81 +/- 6.8 | 22.1 +/- 15.9 | 7.1 +/- 0.7 |

**Table 2.** List of antibodies and immunostaining methods.

| Antigen | Antibody | Dilution | Antigen Retrieval | IHC Staining Kit |
| --- | --- | --- | --- | --- |
| **Amyloid-β** | 4G8<br>BioLegend<br>(800702) | 1:15,000 | Citrate<br>Buffer, in<br>Steamer | Supersensitive Kit |
| **pTau** | AT8<br>Invitrogen<br>(MN1020) | 1:1,600 | Citrate<br>Buffer, in<br>Steamer | Supersensitive Kit |
| **GPNMB** | R&D<br>(AF2550) | 1:500 | Citrate<br>Buffer, in<br>Steamer | ImmPRESS Kit |

Gene regulatory networks were built using pySCENIC (0.10.3) ^61 62^ package default parameters. The inputs for the pySCENIC gene regulatory network analyses were the same filtered expression matrices as for the MEGENA gene co-expression module analyses. Correlations between a list of 1390 human transcription factors (TFs) curated by Lambert et al (2018)^63^ and the genes in the expression matrix were evaluated and co-expression modules with a minimum size of 20 genes were defined. From these, regulons (gene modules sharing a common association with a TF) were built after removing the genes without a recognition motif (based on the hg19-500bp-upstream-7species.mc9nr and hg19-tss-centered-10kb-7species.mc9nr databases provided in the pySCENIC package) for the correlated TF. Only regulons with activator-TFs were retained^61^. Regulons including 50 or more genes were retained for downstream analyses and Cytoscape software (Mac OS version 3.8.0)^60^ was used for their graphical representation.

To evaluate module and regulon enrichment with AD pathology, the AUCell scores for each gene co-expression module or regulon in each nucleus were calculated (aucMaxRank set to 5% of the number of input genes). The statistical analysis was performed using the limma package in R^64^, using the default configuration and the following linear model: ∼pathology+nFeature+pc_mito, where pathology is the average immunohistochemistry quantification value for Aβ or pTau, nFeatures is the total number of distinct features expressed in each nucleus (to account for the fact that nuclei that express a higher number of features may have higher AUCell scores) and pc_mito is the percentage of counts mapping to mitochondrial genes. We also corrected for a potential pseudoreplication bias^65^, by using the duplicateCorrelation function of the limma package with the sample as the “blocking” variable.

Sub-cluster-specificity of modules and regulons were estimated using the regulon_specificity_scores function in pySCENIC^66^. Briefly, the module/regulon specificity score employs the Jensen-Shannon Divergence, a metric previously used to assess cell type specificity of transcripts^67^ and regulons^66^. Modules and regulons with the highest specificity score may be considered subcluster-specific. A specificity score of 1 indicates a gene set that is only expressed in one sub-cluster while a specificity score of 0 indicates an evenly expressed gene set across all sub-clusters.

## Acknowledgements

We thank the donors and their families for the use of human brain tissue in this study and the London Neurodegeneration and Parkinson’s UK Brain Bank staff for making it available. Infrastructure, including particularly the LMS/NIHR Imperial Biomedical Research Centre Flow Cytometry Facility and the Imperial BRC Genomics Facility, were supported by the National Institute for Health Research (NIHR) Biomedical Research Centre (BRC). ST was supported by an “Early Postdoc.Mobility” scholarship (P2GEP3_191446) from the Swiss National Science Foundation and a “Clinical Medicine Plus” scholarship from the Prof Dr Max Cloëtta Foundation (Zurich, Switzerland). PMM acknowledges generous personal support from the Edmond J Safra Foundation and Lily Safra and an NIHR Senior Investigator Award. This work was supported by the UK Dementia Research Institute, which receives its funding from UK DRI Ltd., funded by the UK Medical Research Council, Alzheimer’s Society and Alzheimer’s Research UK.

## Author Contributions

Amy Smith – experimental design, performed experiments, data analysis, interpretation of results, wrote the paper

Karen Davey – performed experiments, data analysis, interpretation of results

Stergios Tsartsalis – co-expression data analysis and visualisation, interpretation of results

Combiz Khozoie – bioinformatic data analysis, differential expression analysis

Nurun Fancy – enrichment analysis and visualisation, interpretation of results

Elijah See – performed immunohistochemistry experiments and image analysis

Eirini Liaptsi – performed immunohistochemistry experiments and image analysis

Maria Weinert – performed optimisation experiments

Aisling McGarry - performed immunohistochemistry experiments

Callum Muirhead – performed immunohistochemistry experiments

Steve Gentleman – cohort selection and neuropathology

David Owen – experimental design, interpretation of results

Paul M. Matthews-experimental design, interpretation of results, wrote the paper

All of the co-authors reviewed and approved the manuscript.

